# Anisotropic Thermal Conductivity in Topologically Networked Protein-MXene Composites

**DOI:** 10.64898/2026.07.10.737764

**Authors:** Mert Vural, Huihun Jung, John A. Tomko, Patrick Hopkins, Melik Demirel

## Abstract

Governing thermal transport in engineered materials creates opportunities to redirect and recover the excess heat generated in electronic and energy-conversion devices. Materials that pair low cross-plane thermal conductivity with high in-plane thermal conductivity are particularly valuable because they confine heat and channel it away from sensitive regions, preventing localized device failure. Two-dimensional crystals are efficient building blocks for such anisotropic thermal conductors, but they are brittle, and the polymer composites used to toughen them usually forfeit much of the intrinsic anisotropy: in conventional percolation-based design, filler fraction is the only handle available, and it governs both in-plane and cross-plane conduction. Here we report a composite of Ti_3_C_2_T_x_ (MXene) nanosheets and squid ring teeth (SRT) inspired recombinant tandem-repeat (TR) proteins in which the protein serves as a molecular template and bridge, setting the spacing between nanosheets with angstrom-level precision through the number of tandem-repeat units and independently of the filler fraction. This structural handle provides a second, independent design parameter. At a fixed MXene loading, the number of repeats tunes the cross-plane conductivity (0.30 to 0.93 W/mK) and, with it, the thermal anisotropy ratio over a wide range (from about 70 down to 17), while the in-plane conductivity stays high (16 to 21 W/mK). We rationalize these trends with a Gaussian Network Model (GNM) of the protein embedded in a two-phase layered medium, which reproduces the measured directional conductivities from a single structural parameter and identifies the protein gallery as the cross-plane bottleneck. Extending the model to a mechanically loaded five-period stack, we find that the anisotropy is robust to reversible compression and twist, changing by only a few percent, so the number of tandem repeats, not the applied strain, is the dominant design handle. Because anisotropy is tuned structurally rather than volumetrically, these protein-MXene composites decouple thermal anisotropy from filler content, pointing toward flexible thermal materials that are not bound by the rules of mixture and percolation.

## INTRODUCTION

The ability to govern thermal transport in materials can offer new possibilities for harvesting and recovering excess thermal energy generated during chemical and physical processes in engineering systems and devices. For instance, heat generated in electronic devices can be confined in a specific geometry and then dissipated away from the devices using an engineered material with anisotropic thermal transport. ^1–6^ Thermal materials with low cross-plane thermal conductivity ensure that the wasted thermal energy remains within the material, and high in-plane thermal conductivity is essential for directing this energy away from operational devices, thereby preventing device failure originating from localized heat.^7^

Since the demonstration of graphene, the family of 2D crystals has expanded continuously and has proven to be an efficient building block for electronic and thermal materials with anisotropic transport characteristics.^1–3,6,8^ Among these 2D crystals, MXenes (Ti3C2Tx and related transition-metal carbides) have emerged as a particularly versatile platform for thermal management, uniquely combining metallic in-plane electrical and thermal conductivity with a chemically tunable, functional-group-terminated surface that can be leveraged to engineer interfacial phonon transport.^41,42^ However, these materials lack the mechanical toughness and flexibility that are essential for propitious applications such as thermal management in smart textiles,^9^ wearables,^10^ and flexible/stretchable electronic devices.^11^ Incorporating polymeric materials into these novel thermal materials is a common approach to circumvent their brittle nature.^12–20^ But it is important to note that the resulting composites could not fully inherit the anisotropic thermal transport mechanism observed in thermal materials constructed solely from 2D crystals.^12–20^ The fundamental limitations in composite systems arise from two distinct mechanisms that influence thermal transport. Thermal transport processes in composite materials that have a filler fraction below the percolation threshold are limited by interfacial effects and thermal boundary resistances^21^, which hinder phonon-mediated thermal transport.^22^ It is possible to mediate this adverse effect by modifying the polymer or surface chemistry of 2D crystals.^23–25^ For example, facilitating a covalent or non-covalent bond between the polymer matrix and 2D crystals can help reduce this thermal boundary resistance.^26^ In these composite systems, thermally conductive fillers based on 2D crystals act more like a doping site, hence resulting in composites with low thermal conductivity and limited thermal anisotropy.^27^ Thermal transport in composite materials with filler fractions exceeding the percolation threshold also suffers from interfacial effects.^12–20,28^ But, the major limitation in these denser thermal composites originates from the percolative distribution of thermally conductive 2D crystals in a polymer matrix.^27,29^ The most promising solution for this problem is governing the organization of 2D crystals in these composite materials.^27^ Engineering the distribution of thermally conductive 2D crystals also helps promote thermal anisotropy in these composites. The orientation of 2D crystals in these composites can be architected through physical (magnetic, pneumatic) forces or a macroscopic template to promote thermal conductivity in a certain direction while hindering thermal conductivity across another.^13,14,16,30–32^ Employing polymer matrices as a molecular template can also result in the alignment of 2D crystals in a certain direction, but these templates cannot alter the separation between 2D crystals parametrically. Consequently, thermal transport in these composites is still bound by the rules of percolation theory. In conventional percolation, the thermal conductivity and thermal anisotropy in composites can be altered by a single parameter: the filler fraction.^27^ Although increasing filler fraction can drastically increase thermal conductivity in composites, it exhibits a more complex relation with thermal anisotropy. For filler fractions slightly above the percolation threshold, increasing filler fraction can drastically improve thermal anisotropy, due to a rapid increase in in-plane thermal conductivity with the addition of nanosheet filler materials.^27,33^ However, once the composite system becomes volumetrically limited, the closing gap between nanosheets increases cross-plane thermal conductivity at a rate that exceeds the increase in in-plane thermal conductivity, thereby leading to a sharp decrease in thermal anisotropy.^34^ A physical system that provides a secondary parameter to adjust thermal conductivity and thermal anisotropy can help mediate this trade-off in composites with filler fractions exceeding the percolation threshold.

Here, we report a composite material system comprising Ti_3_C_2_T_x_ (MXene) nanosheets and synthetic proteins derived from squid ring teeth, which serve as an anisotropic thermal conductor. ^43^ MXene nanosheet fillers with high electrical conductivity can facilitate high in-plane thermal conductivity, while synthetic proteins act as molecular templates and bridges for MXene nanosheets to orchestrate their orientation and govern cross-plane thermal conductivity. The key difference between these synthetic proteins and existing molecular templates used in thermal composites can be stated as their ability to alter interlayer spacing between nanosheets with angstrom-level precision without changing the filler fraction.^35,36^ These synthetic proteins have an amino acid sequence that is composed of tandem repeat (TR) units, each of which possesses specific sequences having a critical role in their assembly characteristics with nanosheets.^35–37^ These assembly characteristics manifest themselves in the structure of these composite materials as a distinct separation distance between nanosheets, which depends on the number of TR units. This structural impact controlled through the number of TR units is reflected in the thermal transport properties of these composites and presents itself as a secondary parameter to adjust thermal conductivity and thermal anisotropy in these composite materials, in complement to filler fraction. The thermal anisotropy and thermal conductivity of these composites can be controlled parametrically without changing the filler material or filler fraction. Hence, it is possible engineer composites from MXene nanosheets with the same filler fraction that exhibit a thermal anisotropy ratio ranging from 17 to 70. Similarly, the thermal conductivity of these composites with the same MXene filler fraction ranges between 16 and 21 W m⁻¹ K⁻¹ in-plane. The structural influence of TR synthetic proteins yields a composite system in which materials with higher thermal anisotropy also exhibit higher thermal conductivity, unlike composites tuned via filler fraction. The intrinsic thermal conductivity of TR proteins, which scales with the number of TR units, also contributes greatly to this phenomenon.^38^ This scaling effect is particularly important for demonstrating composite material systems that can combine high thermal anisotropy with high thermal conductivity without being constrained by the rules of mixtures and percolation theory.

## RESULTS

### Fabrication and processing of MXene/TR composites

We assembled the composite from two building blocks: Ti_3_C_2_T_x_ MXene nanosheets, exfoliated as single- or few-layer sheets from Ti_3_AlC_2_ (MAX phase), and squid-inspired tandem-repeat (TR) proteins (Figure 1a-b). The hydroxyl- and fluorine-terminated surface of the MXene nanosheets supports hydrogen bonding as well as hydrophilic and hydrophobic interactions with the protein matrix (Figure 1a).^36^ These interfacial interactions help minimize phonon scattering at the MXene–protein boundary and thus preserve thermal transport across the composite.^21,22^ We expressed TR proteins in two sizes—corresponding to 4 and 7 tandem-repeat units (TR-n4, TR-n7)—using a rolling-circle amplification strategy that encodes tandem-repeat sequences in a single cloning step.^44^ Each repeat contains one sequence that forms cross-linked β-sheet crystals (PAAASVSTVHHP) and one that forms amorphous domains (YGYGGLYGGLYGGLGY), together with short flanking segments required for amplification (Figure 1b).

**Figure 1.**
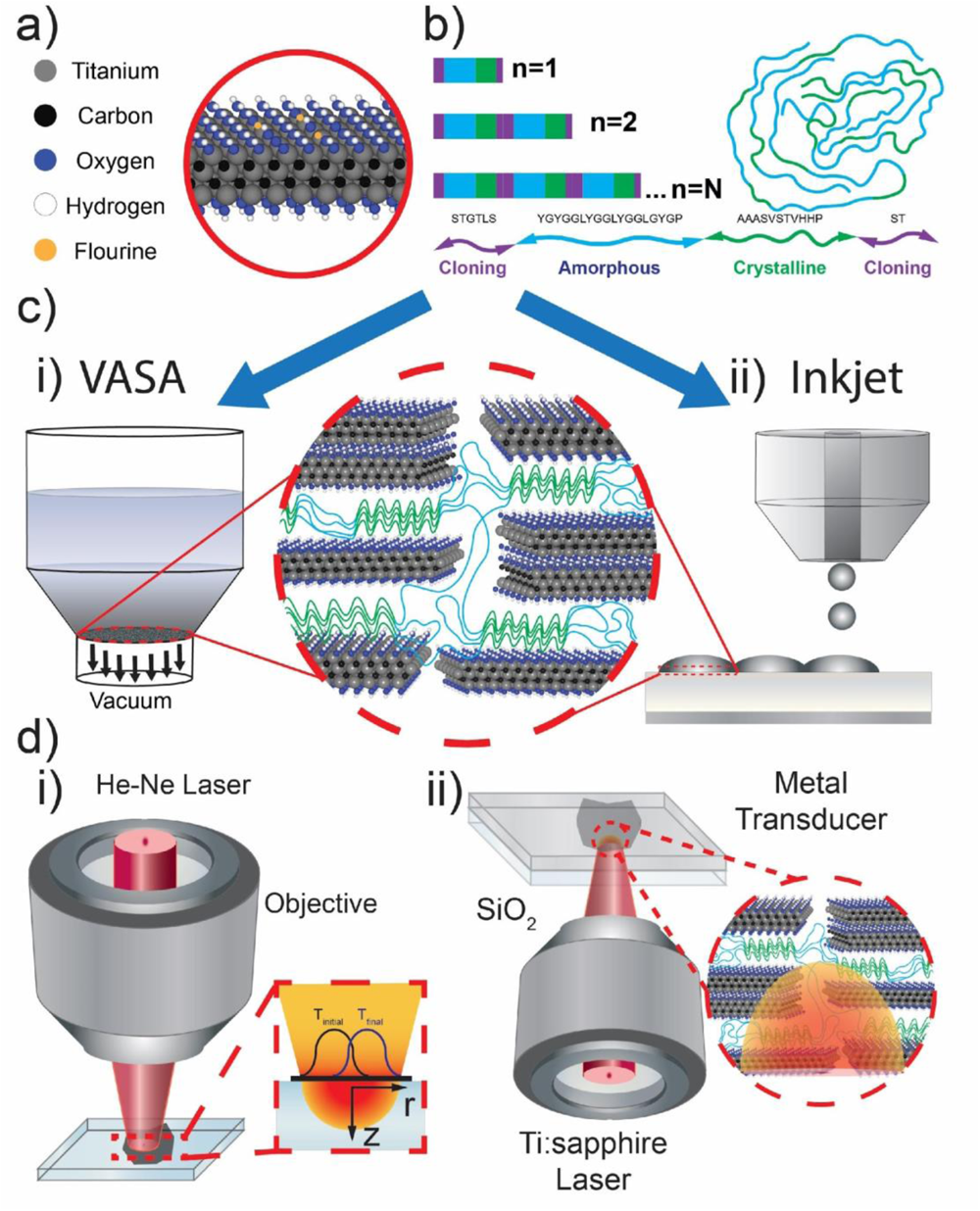
Schematic illustration of a) MXene nanosheets and b) tandem repeat proteins. c) Schematic illustration of fabrication protocols including (i) vacuum-assisted self-assembly (VASA) and (ii) inkjet printing. d) Illustrations of characterization methods of (i) micro-Raman and (ii) TDTR performed on composites fabricated by VASA and inkjet printing, respectively.

To form composite films, we used two complementary solution-processing routes (Figure 1c): vacuum-assisted self-assembly (VASA) and inkjet printing. VASA yields thicker, freestanding films that are well-suited to micro-Raman measurements used to probe bulk (in-plane) thermal transport (Figure 1d(i)).^39^ For VASA, MXene, and TR proteins were dissolved separately (1 and 7.5 mg/mL), mixed, homogenized by stirring and bath sonication, and vacuum-filtered into free-standing films. Inkjet printing instead deposits thinner films on metal-coated glass—the geometry required for time-domain thermoreflectance (TDTR), which is sensitive to cross-plane thermal conductivity (Figure 1d(ii)).^38^ Printing inks used more dilute protein (2 mg/mL) and more concentrated MXene (2.5 mg/mL) DMSO solutions, homogenized as above and jetted onto aluminum-coated glass. Throughout, we distinguish free-standing (VASA) films from printed composite films.

### Structural characterization by X-ray diffraction

To determine how the TR proteins reorganize the MXene nanosheets, we characterized both film types by X-ray diffraction (Figure 2). In free-standing MXene/TR films, the MXene (002) reflection resolves into components corresponding to intercalated water, DMSO, and protein between the nanosheets (Figure 2a). Deconvolving the (002) reflection shows that proteins with different repeat numbers yield distinct nanosheet separations: the protein-intercalation peak shifts to a lower angle—i.e., larger spacing—as the number of repeat units increases (Figure 2a). Printed composite films reproduce the same set of reflections (Figure 2b), although the TR(100) β-sheet peak is weaker and the water peak is broader and centered at a higher angle. We attribute the latter to substrate heating during printing, which limits moisture uptake by the hygroscopic DMSO, protein, and MXene. Extracting interlayer separations confirms that the two routes are structurally equivalent: DMSO-mediated spacings agree to within ±0.3 Å, and protein-mediated spacings exhibit the same repeat-number dependence across both film types (Figure 2c,d). The structural identity imparted by TR-protein assembly is therefore preserved regardless of processing method.

**Figure 2.**
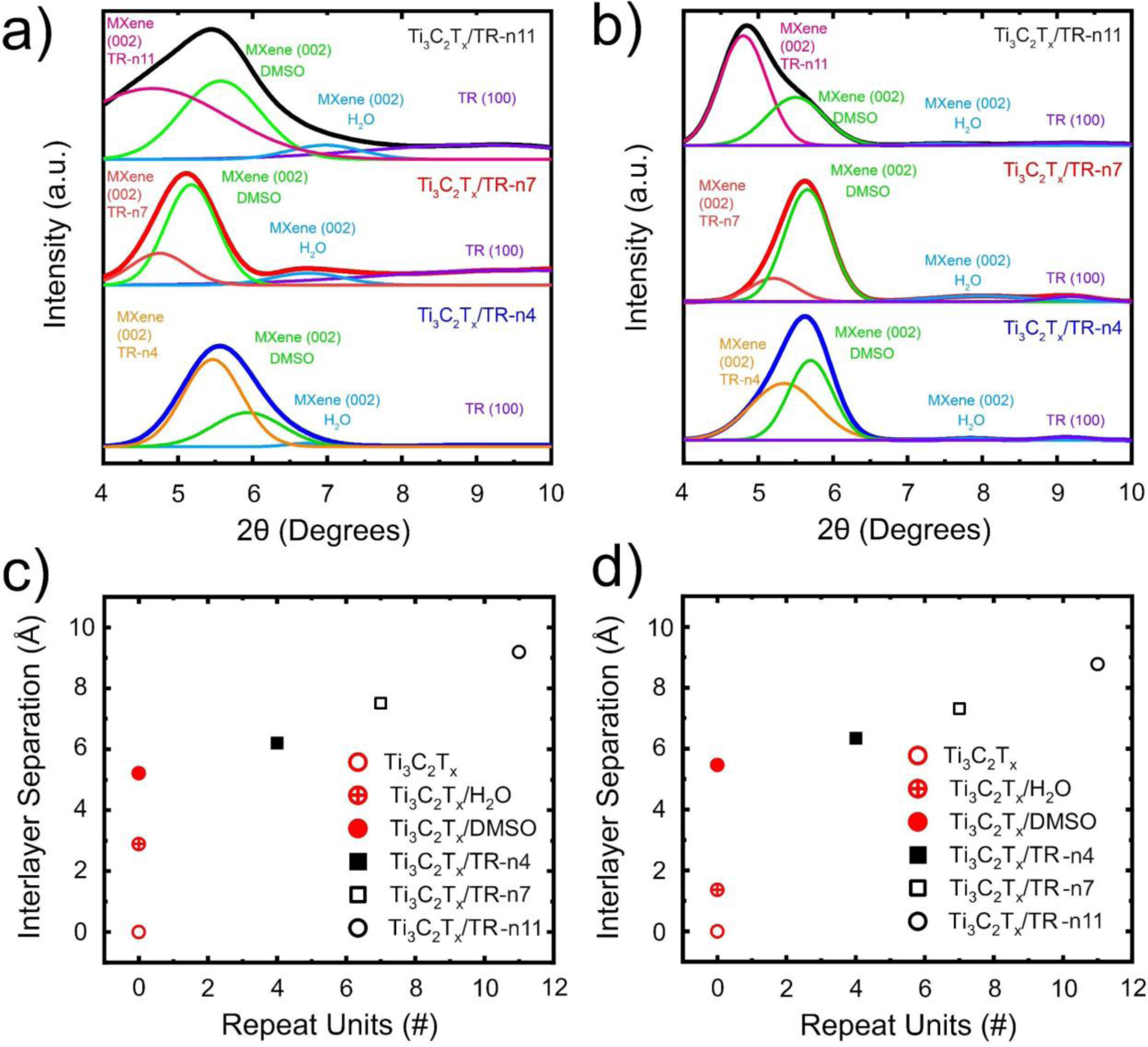
X-ray diffraction patterns of Ti_3_C_2_T_x_ MXene/TR-n4,7 composites fabricated using a) VASA and b) inkjet printing. Intersheet separations stemming from various intercalants in composites of Ti_3_C_2_T_x_ MXene/TR-n4,7 fabricated using c) VASA and d) inkjet printing.

### Composition and filler fraction by thermogravimetric analysis

Because our central claim is that the TR proteins tune thermal transport without altering the filler fraction, we used thermogravimetric analysis (TGA) to confirm that all composites contain essentially the same MXene fraction (Figure 3). Independent of repeat number and processing route, the mass-loss curves are nearly identical (Figure 3a,c), indicating comparable filler content. The derivative curves (Figure 3b,d) resolve the individual components: a small water loss near 100 °C (≤3% v/v in all samples), DMSO evaporation near 190 °C, and protein decomposition between roughly 250 and 350 °C, while MXene remains stable to 500 °C under inert atmosphere. Free-standing films retain slightly more DMSO than printed films (12.5±2% vs 8±2% v/v), consistent with the substrate heating noted above. After excluding solvent, the MXene filler fraction is 26±2% v/v for free-standing films—in agreement with our earlier VASA composites^36,40^—and 24±2% v/v for printed films. The two routes thus yield composites of matched structural and compositional identity, underscoring the templating role of the TR proteins.

**Figure 3.**
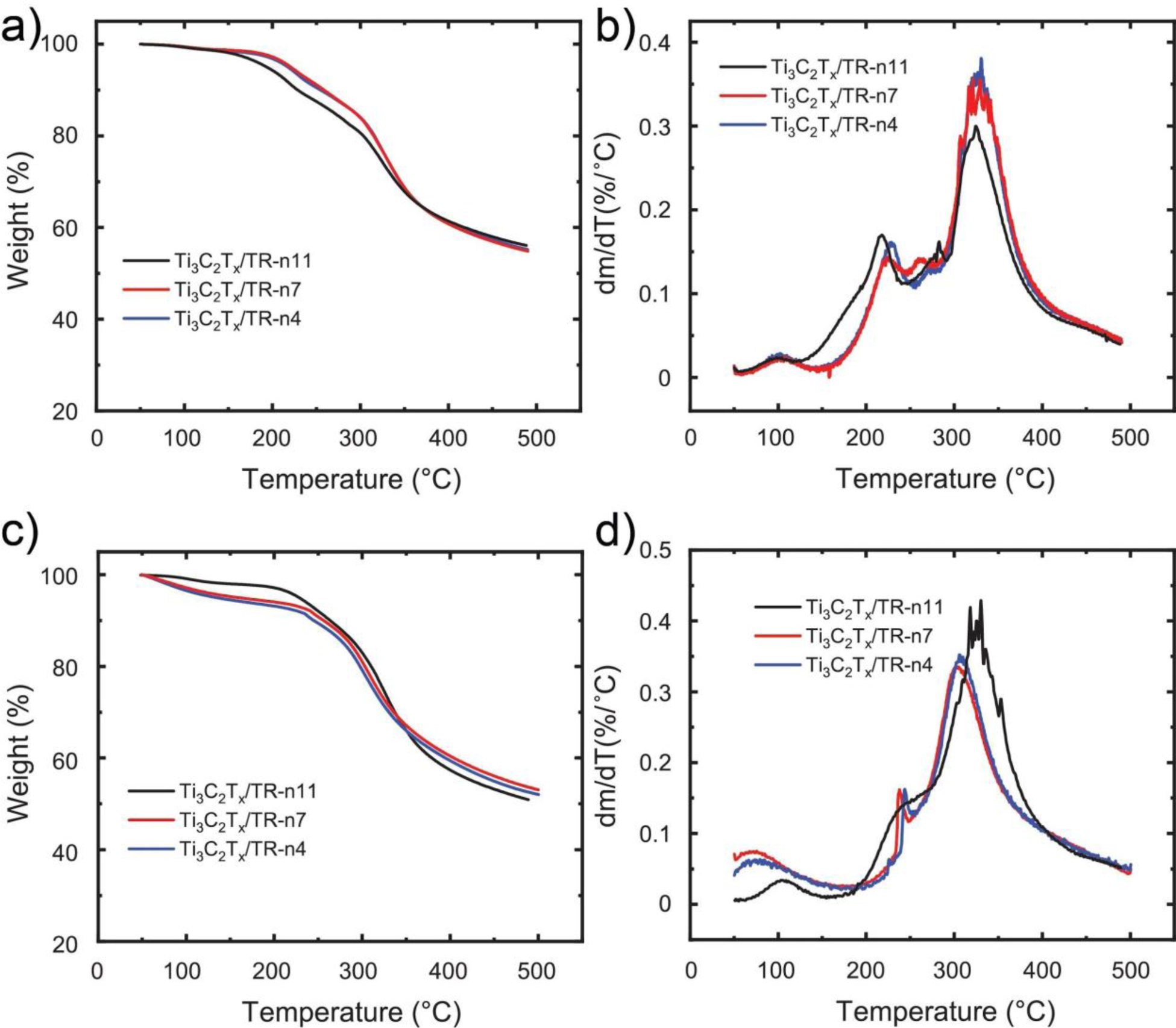
Thermogravimetric analysis (TGA) data shown as a) weight % and b) derivative of weight % for Ti3C2Tx MXene/TR-n4,7 composites fabricated with VASA, and c) weight % and d) derivative of weight % for the same composites fabricated with inkjet-printing.

### Anisotropic thermal transport

To experimentally evaluate the bulk thermal conductivity of this network-structured architecture assembled from TR proteins, we performed micro-Raman measurements on freestanding MXene/TR composites. ^39^ Raman is an effective tool for performing steady-state thermal measurements. The probing laser in Raman spectroscopy can serve as a localized heating source on the sample, and the Raman signal from inelastic phonon scattering can be used to determine the temperature rise at the heated spot.^39^

To gain further insight into the nanoscale heat-transport mechanisms contributing to the high thermal-conductivity anisotropy in our composites, we model the in-plane and cross-plane thermal conductivities using a thermal-resistor approach. The composite is treated as a lamellar stack of two phases: conductive Ti₃C₂Tₓ MXene sheets (intrinsic in-plane conductivity *κ_M_∥*, cross-plane *κ_M⊥_*) separated by intercalated TR-protein galleries (conductivity *κ_p_*). Two structural facts from the composite paper anchor the geometry. First, thermogravimetry shows the MXene volume fraction is essentially constant, *φ_M_* ≈ 0.24–0.26, across n4 and n7 — the filler fraction is *not* the tuning variable. Second, the XRD (002) reflection shifts to a lower angle with *n*, so the protein-gallery height *d_p_*(*n*) grows monotonically with repeat number. The repeat number therefore acts as a secondary structural control parameter — it sets the gallery spacing and the gallery conductivity without changing the amount of MXene present.

The two principal directions correspond to the two classical bounds of a layered medium: in-plane heat runs *along* the sheets (a parallel/arithmetic average dominated by MXene), whereas cross-plane heat must cross the protein galleries *in series* (a harmonic average dominated by the least-conductive phase, the protein).

### In-plane (parallel limit)

Heat conduction along the lamellae adds in parallel, weighting each phase by its volume fraction:

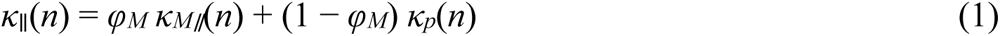

This term is dominated by the MXene network. As the galleries widen with *n*, sheet–sheet registry and overlap of the conductive network weaken, giving a mild monotonic knockdown of the effective in-plane Mxene conductivity, *κ_M∥_*(*n*) = *κ*_M∥,0_ [1 – *η* (*d_p_*(*n*) – *d_p_*(4))], which captures the slow fall of the measured in-plane values (20.7 → 18.8 → 16.1 W m^−1^K^−1^).

### Cross-plane (series limit)

Heat crossing the stack sees the phases in series, so resistances (inverse conductivities) add:

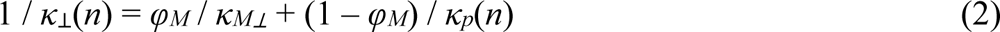

Because *κ_p_* ≪ *κ_M⊥_*, the protein term dominates the sum: the protein gallery is the cross-plane bottleneck and *κ*_⊥_ ≈ *κ_p_*(*n*) / (1 − *φ_M_*). The cross-plane conductivity is therefore a near-direct readout of the protein-gallery conductivity, and inherits whatever *n*-dependence *κ_p_* carries.

Turning to the measured thermal conductivities (Figure 4), we find that as the number of tandem repeats increases—widening the interlayer separation resolved by XRD—the cross-plane conductivity rises steadily, from 0.30 W m^−1^ K^−1^ for pristine MXene to 0.39 for MXene/TR-n4 and 0.93 for MXene/TR-n7 (Figure 4b). Over the same series the in-plane conductivity decreases more gradually, from 20.67 to 18.76 to 16.11 W m^−1^ K^−1^ (Figure 4c).

**Figure 4.**
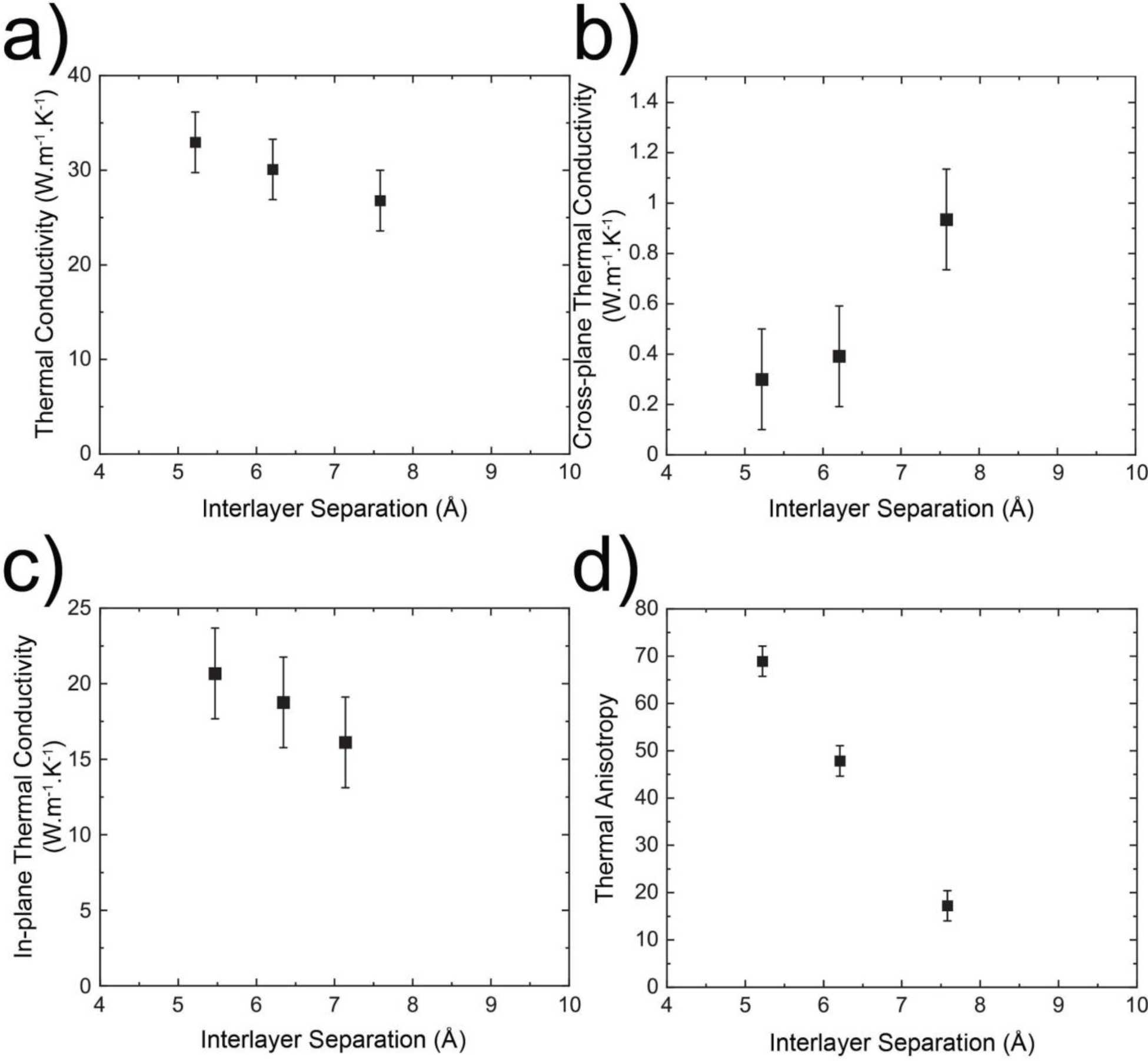
a) Bulk, b) cross-plane, c) in-plane thermal conductivity values of pristine Ti_3_C_2_T_x_ and Ti_3_C_2_T_x_ MXene/TR-n4,7 composites as a function of interlayer separation. d) Thermal anisotropy ratio of pristine Ti_3_C_2_T_x_ and Ti_3_C_2_T_x_ MXene/TR-n4,7 composites as a function of interlayer separation.

These opposing trends set the thermal anisotropy ratio (in-plane divided by cross-plane), shown in Figure 4d. Pristine MXene is the most anisotropic (68.9); inserting progressively longer proteins lowers the ratio to 48.1 for TR-n4 and 17.1 for TR-n7. Because the MXene filler fraction is held constant across the series (Figure 3), the number of tandem repeats serves as a second structural handle on anisotropy, independent of filler loading.

The resistor-network model captures both directional trends with a single structural variable (Figure 5). Fixing the MXene volume fraction at *φ*_M_ ≈ 0.26 and taking the protein-gallery conductivity to scale with the entropic-elasticity tie-chain density, *κ*_p_(*n*) = *κ*_p,0_ + *κ*_p,tie_*ε*_eff_ with *ε*_eff_ = 1 − *β*/*n* and *β* = 4 strands, reproduces the measured in-plane and cross-plane conductivities and the associated anisotropy at *n* = 4 and 7 (Figure 5a,b). The same tie-chain density that governs the mechanical stiffness of these proteins therefore also sets the cross-plane heat flow: as *n* increases, a denser network of heat-carrying tie chains forms within the interlamellar galleries, raising *κ*_p_ (Figure 5c). Notably, the interlayer separation and the tie-chain density are distinct structural channels — the spacing *d*(*n*) grows linearly with repeat number while *ε*_eff_ saturates toward unity (Figure 5d) — so the steep rise in cross-plane conductivity cannot be reproduced by a spacing-driven argument alone but follows naturally from the tie-chain scaling. This modeling framework thus links the thermal anisotropy of the composite directly to the molecular architecture of the protein template.

**Figure 5.**
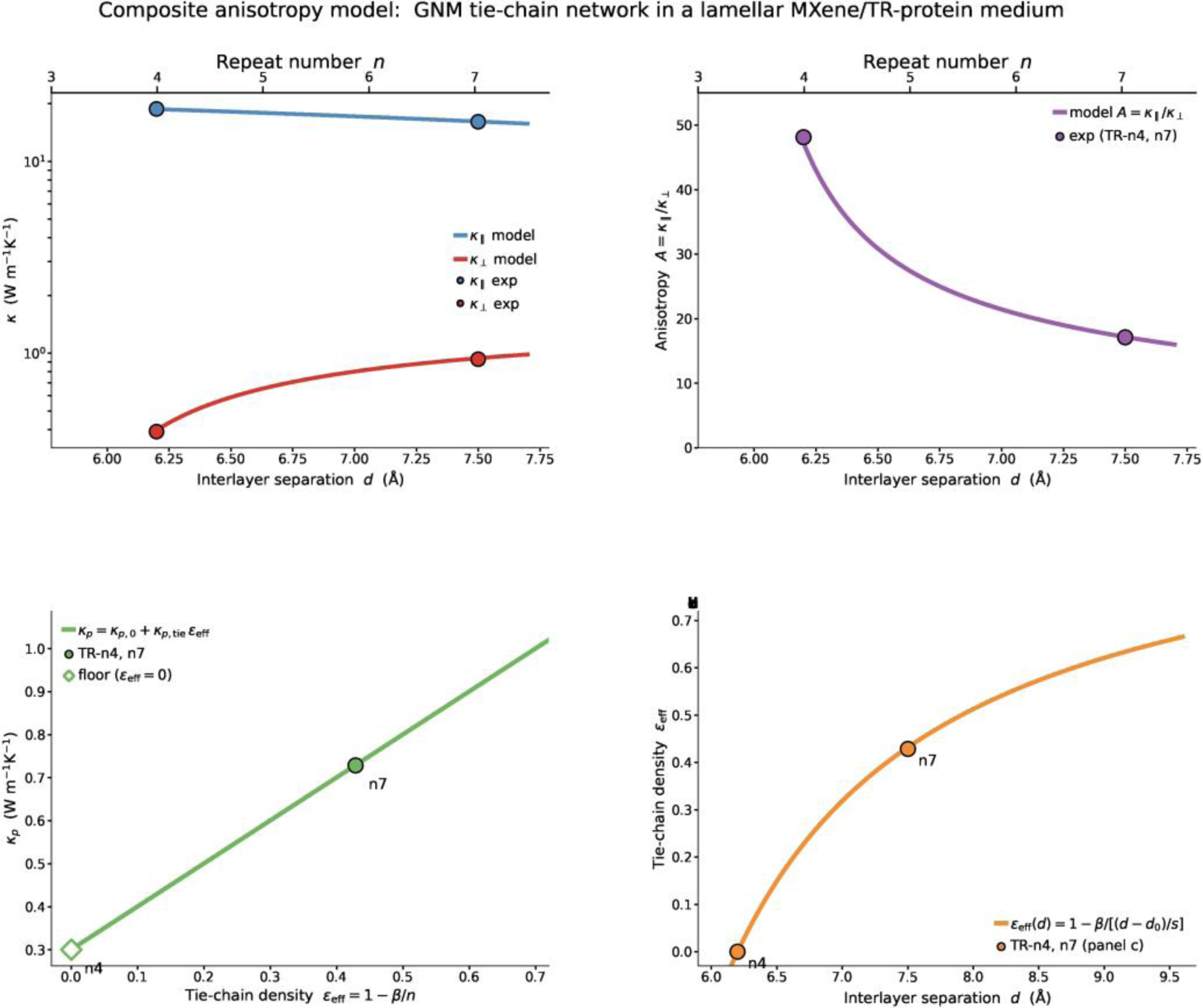
Resistor-network model of anisotropic thermal transport in MXene/TR composites. (a) In-plane (*κ*_∥_, blue) and cross-plane (*κ*_⊥_, red) thermal conductivity versus interlayer separation *d* (lower axis) and the corresponding tandem-repeat number *n* (upper axis); solid curves are the model (Eqs. 1–2), filled symbols are the measured TR-n4 and TR-n7 composites. As *n* increases, *κ*_∥_ falls weakly through dilution and decoupling of the MXene network, while *κ*_⊥_ rises as the protein galleries are enriched with heat-carrying tie chains. (b) The resulting thermal anisotropy ratio *A* = *κ*_∥_/*κ*_⊥_ collapses with *n* at fixed filler fraction (*φ*_M_ ≈ 0.26). (c) The protein-gallery conductivity *κ*_p_ is linear in the effective tie-chain density *ε*_eff_ = 1 − *β*/*n* (*β* = 4), the structure factor that couples the entropic-elasticity protein model to the composite; the open symbol marks the *ε*_eff_ = 0 floor at *n* = 4. (d) *ε*_eff_ expressed as a function of the measured interlayer separation, *ε*_eff_(*d*) = 1 − *β*/[(*d* − *d*_0_)/*s*], using the empirical spacing relation *d*(*n*) = *s*·*n* + *d*_0_ (*s* = 0.43 Å per repeat, *d*_0_ = 4.49 Å) obtained from the XRD (002) shift. Interlayer separation and tie-chain density are distinct structural channels: *d*(*n*) grows linearly with repeat number while *ε*_eff_ saturates, so the steep cross-plane response follows from the tie-chain scaling rather than from spacing alone.

Figure 6 situates the MXene/TR composites among other 2D-material composites. When bulk thermal conductivity (Figure 6a) and anisotropy (Figure 6b) are plotted against filler fraction, the MXene/TR system spans a range that percolation-controlled composites reach only by varying filler content. Because anisotropy here is tuned structurally rather than volumetrically, these composites access a region of the anisotropy-versus-conductivity map (Figure 6c) that is otherwise difficult to reach when filler fraction is the sole design parameter.

**Figure 6.**
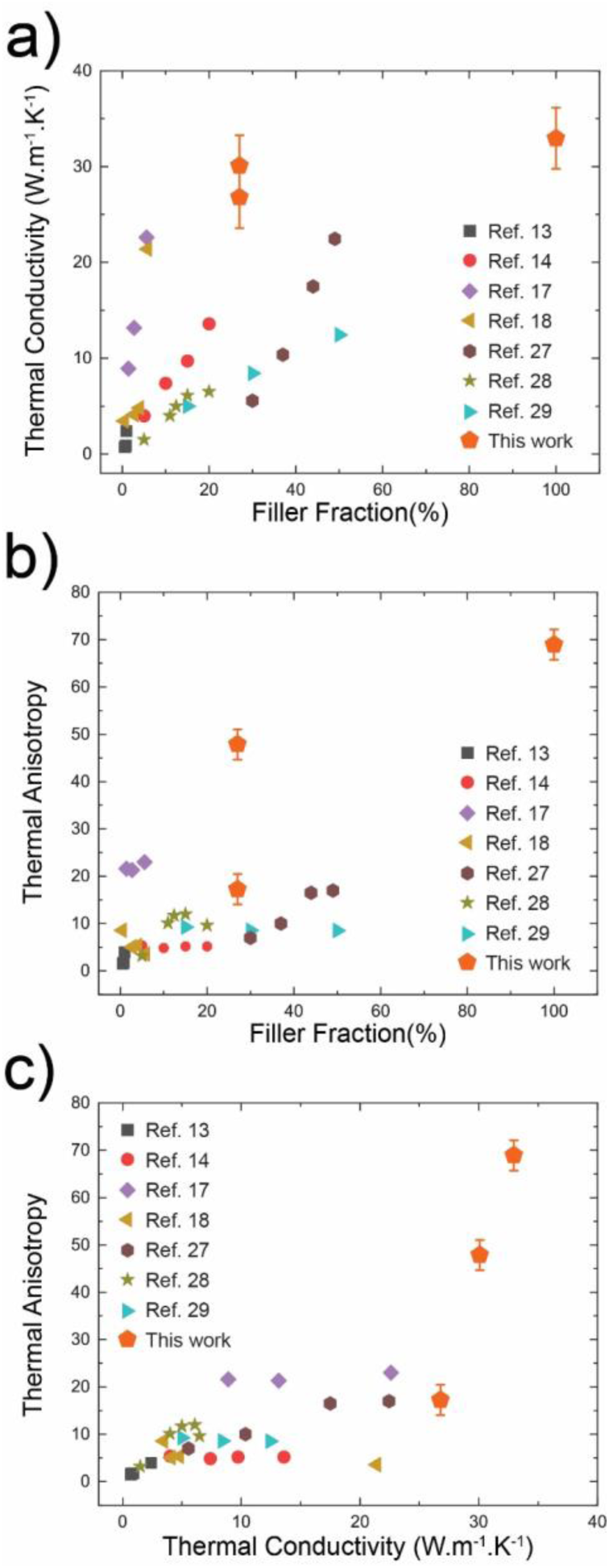
An outlook chart for a) bulk thermal conductivity and b) thermal anisotropy as a function of filler fraction for composites of 2D materials. c) A comparative chart for thermal anisotropy as a function of bulk thermal conductivity of composites composed of 2D materials.

### Mechanical tunability of the anisotropy

Beyond the repeat-number handle, we used the same network/layered-medium framework to examine how the directional response of a stacked composite depends on mechanical load (Figure 7). Modeling a five-period MXene/TR-n4 superlattice (Figure 7a), we relaxed the protein gallery under two modes applied across the stack axial compression and twist about the transport axis — using an anisotropic-network (ANM) description; each mode alters the inter-residue contact lengths and hence the gallery conductivity κp, which we propagate through the series (cross-plane) and parallel (in-plane) averages to obtain the anisotropy A = κ∥/κ⊥. In the undeformed state the composite is strongly anisotropic (κ⊥ = 0.40, κ∥ = 18.7 W m⁻¹ K⁻¹, A = 47), consistent with the measured TR-n4 value.

**Figure 7.**
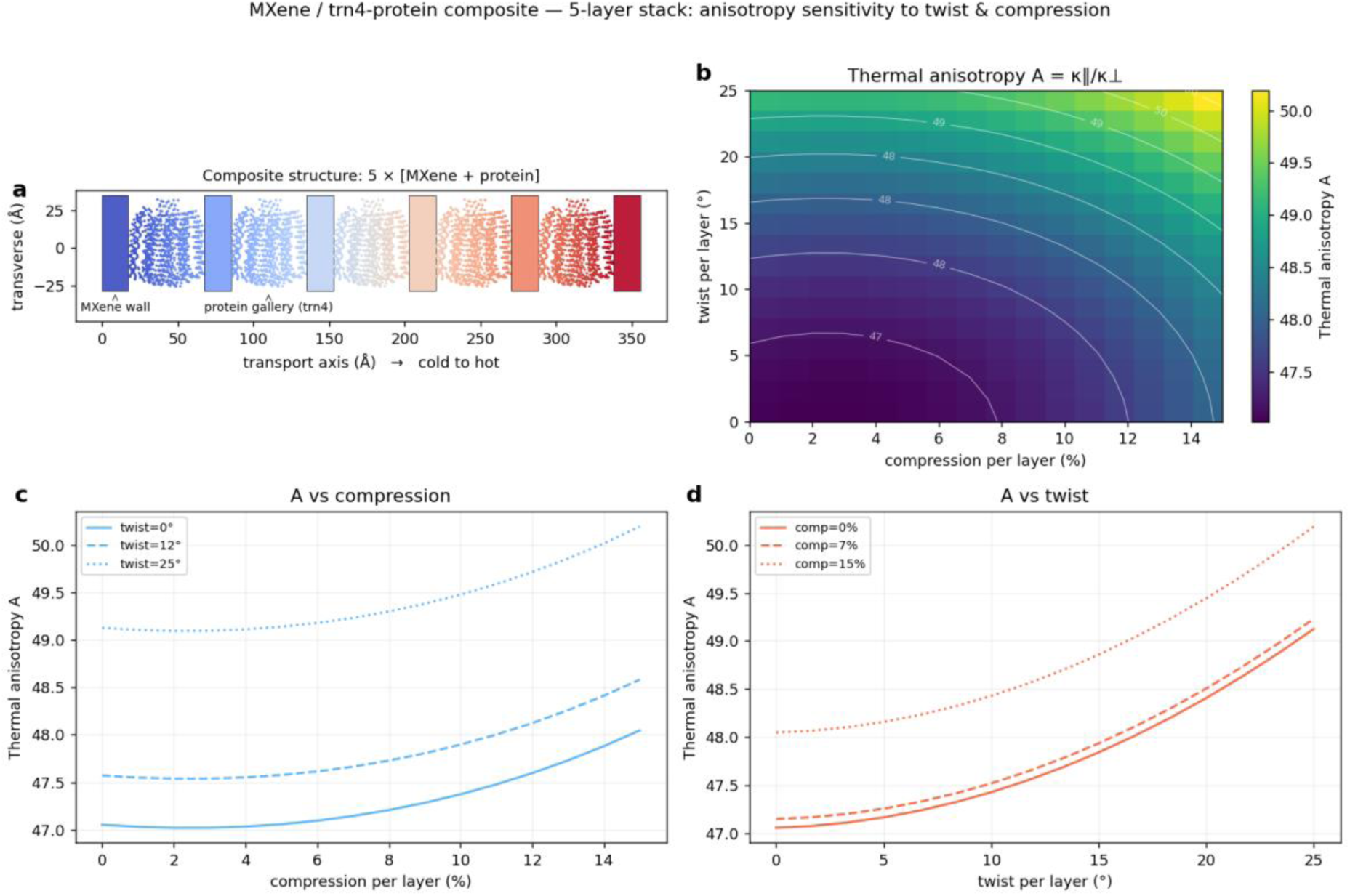
Model prediction of the mechanical sensitivity of the thermal anisotropy of a MXene/TR-protein layered composite. (a) Structure of the modeled composite: five periods of a Ti3C2Tx MXene wall (κ∥ = 71, κ⊥ = 5 W m⁻¹ K⁻¹; wall volume fraction φM = 0.26) alternating with a TR-n4 protein gallery (928 Cα residues) stacked along the transport axis; residues are coloured by the steady-state temperature field, running cold (left) to hot (right). (b) Effective thermal anisotropy A = κ∥/κ⊥ as a function of per-layer compression (0–15 %) and per-layer twist (0–25°); white contours mark integer values of A. (c) A versus compression at fixed twist (0°, 12°, 25°) and (d) A versus twist at fixed compression (0 %, 7 %, 15 %). The gallery conductivity is obtained from the GNM/ANM elastic network (contact cutoff Rc = 10 Å, Grüneisen coupling β = 2), rescaled by the mechanically induced change in network conductance, and combined with the wall through the series (cross-plane) / parallel (in-plane) layered-medium model. For a uniform stack the anisotropy is set by the per-layer deformation and is independent of the number of periods.

Across the full loading range the anisotropy is only weakly tunable (Figure 7b): A varies by at most ≈ 6 % (47 → 50). Twist is the more effective handle, raising A by 4.4 % over 0–25° per layer roughly twice the 2.1 % produced by 0–15 % compression (Figure 7c,d) — and both loads increase A. This follows directly from the layered geometry: because κ∥ ≈ φM κM∥ + (1 − φM) κp is dominated by the MXene sheets it barely responds to the gallery, whereas κ⊥ tracks κp; any deformation that stretches the network and lowers κp therefore lowers κ⊥ faster than κ∥ and raises A. Twist stretches the inter-residue bonds (a quadratic, sign-independent response), whereas compression is nearly flat because axial shortening is largely offset by Poisson lateral expansion. The weak overall response is itself informative: because the anisotropy is fixed by the contrast between the conductive MXene walls and the protein cross-plane bottleneck, it is robust to reversible mechanical perturbation, so the number of tandem repeats — not mechanical load — remains the dominant design parameter.

## DISCUSSION

Taken together, these results show that the number of tandem-repeat units behaves as a structural design parameter that is orthogonal to filler fraction. In a conventional percolation-controlled composite, in-plane and cross-plane conduction are coupled through a single variable—the amount of filler—so that once the system is volumetrically limited, adding filler closes the gap between nanosheets and raises cross-plane conductivity faster than in-plane conductivity, collapsing the anisotropy. Here, the MXene loading is instead held fixed, and the protein directly sets the interlamellar spacing. Lengthening the protein widens this spacing and raises the cross-plane conductivity—consistent with heat conducted through the intercalated protein, whose intrinsic conductivity itself increases with repeat number—while leaving the in-plane MXene network largely intact. The anisotropy ratio therefore falls smoothly from about 70 for pristine MXene to about 17 for MXene/TR-n7, set by molecular design rather than by composition.

This decoupling is what lets the MXene/TR composites reach regions of the anisotropy-versus-conductivity map that are otherwise hard to access when filler fraction is the only handle (Figure 6), while the protein matrix supplies the flexibility and toughness that pristine 2D-crystal films lack. Several questions remain open. Thermal data are reported here for pristine MXene, TR-n4, and TR-n7; extending the measurements to larger repeat numbers would test whether these trends continue as the spacing widens further and whether the cross-plane conductivity eventually saturates. The resistor-network model reproduces the in-plane and cross-plane pathways qualitatively, but a quantitative treatment of the thermal-boundary resistance at the protein–MXene interface would sharpen the mechanistic picture. The residual DMSO and water detected by TGA, and the onset of β-sheet crystallinity at higher repeat numbers, also merit closer study. Even so, the ability to program thermal anisotropy at constant filler content points toward practical, flexible thermal materials for heat management in wearable and stretchable electronics, infrared camouflage, and electromagnetic-interference shielding.

## CONCLUSION

We have shown that synthetic, squid-inspired tandem-repeat proteins can program the anisotropic thermal transport of MXene composites without changing the amount of filler. Using vacuum-assisted self-assembly and inkjet printing, we prepared MXene/TR composites with 4 and 7 repeat units and confirmed, by X-ray diffraction and thermogravimetric analysis, that the two routes yield structurally and compositionally matched films whose only systematic difference is the protein-set spacing between nanosheets. Increasing the number of repeats raised the cross-plane thermal conductivity from 0.30 to 0.93 W m^−1^ K^−1^ while the in-plane conductivity stayed high (16 to 21 W m^−1^ K^−1^), so that the thermal anisotropy ratio could be tuned from roughly 70 to 17 at a fixed MXene loading. A Gaussian/anisotropic-network model of the protein embedded in a two-phase layered medium reproduces both directional conductivities from this single structural parameter and attributes the switch to the widening protein gallery that gates cross-plane transport; extending the model to a mechanically loaded, five-period stack shows that the resulting anisotropy is intrinsic to the layered architecture, changing by only a few percent under reversible compression or twist. Because this control is exercised through molecular design rather than filler fraction — and is preserved under mechanical deformation — the tandem-repeat number provides a second, independent, and mechanically robust handle on thermal transport that conventional rule-of-mixture and percolation frameworks do not offer. Combined with the flexibility of the protein matrix, this strategy offers a route to programmable thermal materials for heat management in flexible and wearable electronics, infrared camouflage, and electromagnetic-interference shielding.

## Acknowledgments

The authors thank staff members of Penn State MRI and Huck User Facilities for helping data collection and sample characterization.

## Funding

M.C.D., S.S., T.H., and H.J. were supported by the Army Research Office (grant no. W911NF-16-1-0019), DARPA (D19AC00016), and the Huck Endowment of Pennsylvania State University. P.H. and J.T. were also supported by DARPA (D19AC00016).

## Author contributions

M.C.D. and P.H. supervised the project. S.S. prepared the composite samples, and worked on characterization. H.J. and T.H. performed the protein expression and purification. J.T. performed the thermal conductivity measurements. All authors participated in manuscript editing, revisions, discussions, and interpretation of the data.

## Competing interests

Authors have pending or existing patent applications

## Data and materials availability

All data is available in the main text or the supplementary materials.

## Materials and Methods

### Synthesis of Tandem Repeat Proteins

The first squid-inspired tandem repeat proteins are obtained by protein expression, gene sequencing, and protein design. A sequence-verified plasmid is transferred to E. coli. After colony inoculation and fermentation, the cell is harvested. The resultant product is purified to get tandem repeat proteins.

### Synthesis of MXene

MXene was produced from the MAX Phase precursor Ti3AlC2 by selective etching. A 6 M 20 mL HCl solution was poured into the Teflon round-bottom flask. 1 g of lithium fluoride and 1 g of MAX-phase raw material are slowly added. They mixed at 60°C for 24 hours. The solution is centrifuged for 30 minutes at 6000 rpm, and the precipitate is washed with deionized water. This is repeated until the pH becomes 6. The resulting solution is centrifuged for 1 hour at 6000 rpm, and the water is exchanged with DMSO, repeated 3 times. MXene in DMSO is made homogeneous by tip sonication, yielding a stable colloidal solution.

### Inkjet printing

2D assembly can be done with 2 methods. For inkjet printing, 0.95 mg/ml TR protein and 2.25 mg/ml Mxene solutions are sonicated. Jetlab 4 (Microfab Technologies Inc., Plano, TX) with an orifice diameter of 120 μm is used. The metal substrates are heated to 80°C to evaporate DMSO during the inkjet printing process. Each inkjet is done for 50 layers of a circle shape.

### Vacuum-assisted self-assembly

Another method is vacuum-assisted self-assembly (VASA). 20 ml 7.5 mg/ml protein-DMSO solution is mixed with 30 ml 1mg/ml MXene-DMSO. The mixture is bath sonicated for 15 minutes. The resulting solution is vacuum-filtered using an Anodisc membrane filter with a diameter of 47 mm and a pore size of 0.2 mm.

### Electron Microscopy

SEM imaging was done using a ZEISS 55 Ultra FESEM at 10 kV voltage.

### X-Ray Diffraction

Wide-angle X-Ray scattering characterization was performed using a SAXS/WAXS system (Xeuss 2.0 HR, Xenocs, France) with a microfocus sealed copper tube under vacuum at room temperature. The wavelength was 1.54 Å (50 kV, 0.6 mA). Pilatus3 R200K detector was utilized. The distance between the sample and the detector was approximately 0.156 m.

### Thermal Conductivity

We measure the thermal conductivity of the MXene/SRT composites with time-domain thermoreflectance (TDTR) and Raman spectroscopy. The TDTR measurements are performed on ink-jet printed films, using a sample geometry and bi-directional thermal analysis identical to that of Ref. [1], while the Raman measurements are performed on free-standing, VASA printed composites. For these VASA films, we first calibrate the thermo-optic coefficient of the MXene/SRT composite by performing temperature-dependent Raman measurements from room temperature to 200 C. We then measure the Raman shift with varying laser power to extract the sample thermal conductivity (*κmeasured*=√*κzκr*). As TDTR is primarily sensitive to the cross-plane thermal conductivity, *κz*, we separate and report the values for in-plane, *κz*, and cross-plane thermal conductivities.

## REFERENCES

(1) Jang, H.; Wood, J. D.; Ryder, C. R.; Hersam, M. C.; Cahill, D. G. Anisotropic Thermal Conductivity of Exfoliated Black Phosphorus. Adv. Mater. 2015, 27 (48), 8017–8022. 10.1002/adma.201503466.

(2) Luo, Z.; Maassen, J.; Deng, Y.; Du, Y.; Garrelts, R. P.; Lundstrom, M. S.; Ye, P. D.; Xu, X. Anisotropic In-Plane Thermal Conductivity Observed in Few-Layer Black Phosphorus. Nat. Commun. 2015, 6 (1), 8572. 10.1038/ncomms9572.

(3) Jang, H.; Ryder, C. R.; Wood, J. D.; Hersam, M. C.; Cahill, D. G. 3D Anisotropic Thermal Conductivity of Exfoliated Rhenium Disulfide. Adv. Mater. 2017, 29 (35), 1700650. 10.1002/adma.201700650.

(4) Sun, B.; Haunschild, G.; Polanco, C.; Ju, J. (Zi-J.; Lindsay, L.; Koblmüller, G.; Koh, Y. K. Dislocation-Induced Thermal Transport Anisotropy in Single-Crystal Group-III Nitride Films. Nat. Mater. 2019, 18 (2), 136–140. 10.1038/s41563-018-0250-y.

(5) Hoque, M. S. Bin; Koh, Y. R.; Braun, J. L.; Mamun, A.; Liu, Z.; Huynh, K.; Liao, M. E.; Hussain, K.; Cheng, Z.; Hoglund, E. R.; Olson, D. H.; Tomko, J. A.; Aryana, K.; Galib, R.; Gaskins, J. T.; Elahi, M. M. M.; Leseman, Z. C.; Howe, J. M.; Luo, T.; Graham, S.; Goorsky, M. S.; Khan, A.; Hopkins, P. E. High In-Plane Thermal Conductivity of Aluminum Nitride Thin Films. ACS Nano 2021, 15 (6), 9588–9599. 10.1021/acsnano.0c09915.

(6) Kim, S. E.; Mujid, F.; Rai, A.; Eriksson, F.; Suh, J.; Poddar, P.; Ray, A.; Park, C.; Fransson, E.; Zhong, Y.; Muller, D. A.; Erhart, P.; Cahill, D. G.; Park, J. Extremely Anisotropic van Der Waals Thermal Conductors. Nature 2021, 597 (7878), 660–665. 10.1038/s41586-021-03867-8.

(7) Pop, E. Energy Dissipation and Transport in Nanoscale Devices. Nano Res. 2010, 3 (3), 147–169. 10.1007/s12274-010-1019-z.

(8) Fu, Y.; Hansson, J.; Liu, Y.; Chen, S.; Zehri, A.; Samani, M. K.; Wang, N.; Ni, Y.; Zhang, Y.; Zhang, Z.-B.; Wang, Q.; Li, M.; Lu, H.; Sledzinska, M.; Torres, C. M. S.; Volz, S.; Balandin, A. A.; Xu, X.; Liu, J. Graphene Related Materials for Thermal Management. 2D Mater. 2019, 7 (1), 12001. 10.1088/2053-1583/ab48d9.

(9) Po-Chun, H.; Y., S. A.; B., C. P.; Chong, L.; Yucan, P.; Jin, X.; Shanhui, F.; Yi, C. Radiative Human Body Cooling by Nanoporous Polyethylene Textile. Science (80-.). 2016, 353 (6303), 1019–1023. 10.1126/science.aaf5471.

(10) Jayathilaka, W. A. D. M.; Qi, K.; Qin, Y.; Chinnappan, A.; Serrano-García, W.; Baskar, C.; Wang, H.; He, J.; Cui, S.; Thomas, S. W.; Ramakrishna, S. Significance of Nanomaterials in Wearables: A Review on Wearable Actuators and Sensors. Adv. Mater. 2019, 31 (7), 1805921. 10.1002/adma.201805921.

(11) Sun, B.; Huang, X. Seeking Advanced Thermal Management for Stretchable Electronics. *npj Flex*. Electron. 2021, 5 (1), 12. 10.1038/s41528-021-00109-9.

(12) Teng, C.-C.; Ma, C.-C. M.; Lu, C.-H.; Yang, S.-Y.; Lee, S.-H.; Hsiao, M.-C.; Yen, M.-Y.; Chiou, K.-C.; Lee, T.-M. Thermal Conductivity and Structure of Non-Covalent Functionalized Graphene/Epoxy Composites. Carbon N. Y. 2011, 49 (15), 5107–5116. 10.1016/j.carbon.2011.06.095.

(13) Kumar, P.; Yu, S.; Shahzad, F.; Hong, S. M.; Kim, Y.-H.; Koo, C. M. Ultrahigh Electrically and Thermally Conductive Self-Aligned Graphene/Polymer Composites Using Large-Area Reduced Graphene Oxides. Carbon N. Y. 2016, 101, 120–128. 10.1016/j.carbon.2016.01.088.

(14) Lian, G.; Tuan, C.-C.; Li, L.; Jiao, S.; Wang, Q.; Moon, K.-S.; Cui, D.; Wong, C.-P. Vertically Aligned and Interconnected Graphene Networks for High Thermal Conductivity of Epoxy Composites with Ultralow Loading. Chem. Mater. 2016, 28 (17), 6096–6104. 10.1021/acs.chemmater.6b01595.

(15) Hamidinejad, S. M.; Chu, R. K. M.; Zhao, B.; Park, C. B.; Filleter, T. Enhanced Thermal Conductivity of Graphene Nanoplatelet–Polymer Nanocomposites Fabricated via Supercritical Fluid-Assisted in Situ Exfoliation. ACS Appl. Mater. Interfaces 2018, 10 (1), 1225–1236. 10.1021/acsami.7b15170.

(16) Song, S. H.; Park, K. H.; Kim, B. H.; Choi, Y. W.; Jun, G. H.; Lee, D. J.; Kong, B.-S.; Paik, K.-W.; Jeon, S. Enhanced Thermal Conductivity of Epoxy–Graphene Composites by Using Non-Oxidized Graphene Flakes with Non-Covalent Functionalization. Adv. Mater. 2013, 25 (5), 732–737. 10.1002/adma.201202736.

(17) Yavari, F.; Fard, H. R.; Pashayi, K.; Rafiee, M. A.; Zamiri, A.; Yu, Z.; Ozisik, R.; Borca-Tasciuc, T.; Koratkar, N. Enhanced Thermal Conductivity in a Nanostructured Phase Change Composite Due to Low Concentration Graphene Additives. J. Phys. Chem. C 2011, 115 (17), 8753–8758. 10.1021/jp200838s.

(18) Yu, A.; Ramesh, P.; Itkis, M. E.; Bekyarova, E.; Haddon, R. C. Graphite Nanoplatelet−Epoxy Composite Thermal Interface Materials. J. Phys. Chem. C 2007, 111 (21), 7565–7569. 10.1021/jp071761s.

(19) Chen, L.; Cao, Y.; Zhang, X.; Guo, X.; Song, P.; Chen, K.; Lin, J. Anisotropic and High Thermal Conductivity of Epoxy Composites Containing Multilayer Ti3C2Tx MXene Nanoflakes. J. Mater. Sci. 2020, 55 (35), 16533–16543. 10.1007/s10853-020-05177-2.

(20) Aakyiir, M.; Araby, S.; Michelmore, A.; Meng, Q.; Amer, Y.; Yao, Y.; Li, M.; Wu, X.; Zhang, L.; Ma, J. Elastomer Nanocomposites Containing MXene for Mechanical Robustness and Electrical and Thermal Conductivity. Nanotechnology 2020, 31 (31), 315715. 10.1088/1361-6528/ab88eb.

(21) Giri, A.; Hopkins, P. E. A Review of Experimental and Computational Advances in Thermal Boundary Conductance and Nanoscale Thermal Transport across Solid Interfaces. Adv. Funct. Mater. 2020, 30 (8), 1903857. 10.1002/adfm.201903857.

(22) Prasher, R. Thermal Interface Materials: Historical Perspective, Status, and Future Directions. Proc. IEEE 2006, 94 (8), 1571–1586. 10.1109/JPROC.2006.879796.

(23) Hopkins, P. E.; Baraket, M.; Barnat, E. V; Beechem, T. E.; Kearney, S. P.; Duda, J. C.; Robinson, J. T.; Walton, S. G. Manipulating Thermal Conductance at Metal–Graphene Contacts via Chemical Functionalization. Nano Lett. 2012, 12 (2), 590–595. 10.1021/nl203060j.

(24) Foley, B. M.; Hernández, S. C.; Duda, J. C.; Robinson, J. T.; Walton, S. G.; Hopkins, P. E. Modifying Surface Energy of Graphene via Plasma-Based Chemical Functionalization to Tune Thermal and Electrical Transport at Metal Interfaces. Nano Lett. 2015, 15 (8), 4876–4882. 10.1021/acs.nanolett.5b00381.

(25) Walton, S. G.; Foley, B. M.; Hernández, S. C.; Boris, D. R.; Baraket, M.; Duda, J. C.; Robinson, J. T.; Hopkins, P. E. Plasma-Based Chemical Functionalization of Graphene to Control the Thermal Transport at Graphene-Metal Interfaces. Surf. Coatings Technol. 2017, 314, 148–154. 10.1016/j.surfcoat.2016.12.085.

(26) Hopkins, P. E. Thermal Transport across Solid Interfaces with Nanoscale Imperfections: Effects of Roughness, Disorder, Dislocations, and Bonding on Thermal Boundary Conductance. ISRN Mech. Eng. 2013, 2013, 682586. 10.1155/2013/682586.

(27) Shtein, M.; Nadiv, R.; Buzaglo, M.; Kahil, K.; Regev, O. Thermally Conductive Graphene-Polymer Composites: Size, Percolation, and Synergy Effects. Chem. Mater. 2015, 27 (6), 2100–2106. 10.1021/cm504550e.

(28) Shahil, K. M. F.; Balandin, A. A. Graphene–Multilayer Graphene Nanocomposites as Highly Efficient Thermal Interface Materials. Nano Lett. 2012, 12 (2), 861–867. 10.1021/nl203906r.

(29) Shtein, M.; Nadiv, R.; Buzaglo, M.; Regev, O. Graphene-Based Hybrid Composites for Efficient Thermal Management of Electronic Devices. ACS Appl. Mater. Interfaces 2015, 7 (42), 23725–23730. 10.1021/acsami.5b07866.

(30) Song, G.; Kang, R.; Guo, L.; Ali, Z.; Chen, X.; Zhang, Z.; Yan, C.; Lin, C.-T.; Jiang, N.; Yu, J. Highly Flexible Few-Layer Ti3C2 MXene/Cellulose Nanofiber Heat-Spreader Films with Enhanced Thermal Conductivity. New J. Chem. 2020, 44 (17), 7186–7193. 10.1039/D0NJ00672F.

(31) Yuan, C.; Duan, B.; Li, L.; Xie, B.; Huang, M.; Luo, X. Thermal Conductivity of Polymer-Based Composites with Magnetic Aligned Hexagonal Boron Nitride Platelets. ACS Appl. Mater. Interfaces 2015, 7 (23), 13000–13006. 10.1021/acsami.5b03007.

(32) Song, N.; Jiao, D.; Cui, S.; Hou, X.; Ding, P.; Shi, L. Highly Anisotropic Thermal Conductivity of Layer-by-Layer Assembled Nanofibrillated Cellulose/Graphene Nanosheets Hybrid Films for Thermal Management. ACS Appl. Mater. Interfaces 2017, 9 (3), 2924–2932. 10.1021/acsami.6b11979.

(33) Yang, W.; Zhao, Z.; Wu, K.; Huang, R.; Liu, T.; Jiang, H.; Chen, F.; Fu, Q. Ultrathin Flexible Reduced Graphene Oxide/Cellulose Nanofiber Composite Films with Strongly Anisotropic Thermal Conductivity and Efficient Electromagnetic Interference Shielding. J. Mater. Chem. C 2017, 5 (15), 3748–3756. 10.1039/C7TC00400A.

(34) Wu, Z.; Xu, C.; Ma, C.; Liu, Z.; Cheng, H.-M.; Ren, W. Synergistic Effect of Aligned Graphene Nanosheets in Graphene Foam for High-Performance Thermally Conductive Composites. Adv. Mater. 2019, 31 (19), 1900199. 10.1002/adma.201900199.

(35) Vural, M.; Lei, Y.; Pena-Francesch, A.; Jung, H.; Allen, B.; Terrones, M.; Demirel, M. C. Programmable Molecular Composites of Tandem Proteins with Graphene Oxide for Efficient Bimorph Actuators. Carbon N. Y. 2017, 118, 404–412. 10.1016/j.carbon.2017.03.053.

(36) Vural, M.; Zhu, H.; Pena-Francesch, A.; Jung, H.; Allen, B. D.; Demirel, M. C. Self-Assembly of Topologically Networked Protein–Ti3C2Tx MXene Composites. ACS Nano 2020, 14 (6), 6956–6967. 10.1021/acsnano.0c01431.

(37) Demirel, M. C.; Vural, M.; Terrones, M. Composites of Proteins and 2D Nanomaterials. Adv. Funct. Mater. 2018, 28 (0), 1704990. 10.1002/adfm.201704990.

(38) Tomko, J. A.; Pena-Francesch, A.; Jung, H.; Tyagi, M.; Allen, B. D.; Demirel, M. C.; Hopkins, P. E. Tunable Thermal Transport and Reversible Thermal Conductivity Switching in Topologically Networked Bio-Inspired Materials. Nat. Nanotechnol. 2018, 13 (10), 959–964. 10.1038/s41565-018-0227-7.

(39) Malekpour, H.; Balandin, A. A. Raman-Based Technique for Measuring Thermal Conductivity of Graphene and Related Materials. J. Raman Spectrosc. 2018, 49 (1), 106–120. 10.1002/jrs.5230.

(40) Vural, M.; Pena-Francesch, A.; Bars-Pomes, J.; Jung, H.; Gudapati, H.; Hatter, C. B.; Allen, B.; Anasori, B.; Ozbolat, I. T.; Gogotsi, Y.; Demirel, M. C. Inkjet Printing of Self-Assembled 2D Titanium Carbide and Protein Electrodes for Stimuli Responsive Electromagnetic Shielding. Adv. Funct. Mater. 2018, 28, 1801972.

(41) Wang, J. F.; Shen, M. M.; Liu, Z. X.; Wang, W. J. MXene Materials for Advanced Thermal Management and Thermal Energy Utilization. Nano Energy 2022, 97, 107177. 10.1016/j.nanoen.2022.107177.

(42) Shahzad, F.; Alhabeb, M.; Hatter, C. B.; Anasori, B.; Hong, S. M.; Koo, C. M.; Gogotsi, Y. Electromagnetic Interference Shielding with 2D Transition Metal Carbides (MXenes). Science 2016, 353 (6304), 1137–1140. 10.1126/science.aag2421.

(43) Pena-Francesch, A.; Demirel, M. C. Squid-Inspired Tandem Repeat Proteins: Functional Fibers and Films. Front. Chem. 2019, 7, 69. 10.3389/fchem.2019.00069.

(44) Amiram, M.; Quiroz, F. G.; Callahan, D. J.; Chilkoti, A. A Highly Parallel Method for Synthesizing DNA Repeats Enables the Discovery of ‘Smart’ Protein Polymers. Nat. Mater. 2011, 10 (2), 141–148. 10.1038/nmat2942.

